# Testing the reality gap with kilobots performing two ant-inspired foraging behaviours

**DOI:** 10.1101/2024.06.02.596655

**Authors:** Niamh Ellis, Alexander Caravaggio, Jonathan Kelly, Ian Young, Fabio Manfredini, Maria Elena Giannaccini

## Abstract

Robotics looks to nature for inspiration to perform effectively in unstructured environments and can be used as a platform to test biological hypotheses. Social animals often share information about food source locations: one example is tandem running in ants, where a leader guides a naive recruit to a known profitable food site. This is extremely advantageous as it allows the sharing of important information among colony members, but it also has costs, such as waiting time inside the nest when no leaders are around, and a reduced walking speed for the tandem couple compared to individual ants. Whether and when these costs outweigh the benefits is not well understood as it is challenging to observe complex social behaviours in nature. We developed two kilobot-based approaches to compare tandem running and lone scout foraging, where an ant searches for food without any previous knowledge of its location: one approach based on real-life experiments and one on computer simulations. We investigated the role that the size of the search arena played in the effectiveness of foraging. Tandem pairs were faster for all three arena sizes; however, this result was reversed in the simulations. These results highlight the inconsistencies between simulation and real-life kilobot experiments, previously reported for other systems and known as reality gap. Further testing is needed to inform on whether robotic applications should utilise agents with the same roles and capabilities for search, detection and repair-type tasks as simulations, or whether instead the two approaches should be treated separately.

## INTRODUCTION

The robotics industry has been fast developing, with a large number of robots being currently employed in areas such as manufacturing [1] or the military: the number of applications is expected to continue its growth as the technology develops. This growth has brought the need to design robots that can function in unstructured and chaotic environments, very different from the highly structured environments that robots have typically been built for [2]. Designing and programming a robot that can respond to the vast and unpredictable possibilities of unstructured environments is extremely challenging. To overcome these challenges researchers have often looked to the natural world to understand how the creatures within behave and use this as an inspiration for robots’ behaviour. At the same time, robots can be used as models of animal systems to test hypotheses regarding their behaviour [3], [4].

A multitude of problems can be solved using a number of simple agents working in parallel rather than a single complex agent: this is identified as swarm robotics [5]. Robot swarms are cheap, re-configurable and function as simple modular units that can tackle a task as a team: for these reasons, swarm robotics has been identified as one of the current high impact challenges in robotics [6]. Systems that implement swarm intelligence possess key properties such as self-organization, scalability, and robustness [7]. The potential applications for swarm robotics are endless, from the medical field with microrobotic swarms that can significantly increase dose delivery [8], to helping humans in social tasks such as brainstorming [9], and underwater exploration [10]. However, it has proven often challenging to predict how robotic swarms behave in an experimental setup using simulations, a step that is needed before robotics can be implemented in real life scenarios: this discrepancy is often referred to as “reality gap” [11]. While it has been suggested that experiments with real robots can be avoided if using a different model for the simulated behaviour [12] [13], this approach might miss important aspects of a real life scenario. While this reality gap has been thoroughly documented for swarm robotics [14], [15], [16] there are no works, to our knowledge, that explored the reality gap using kilobots to compare two different strategies of foraging behaviour.

Kilobots are swarm robots designed by Harvard [17]. Kilobots communicate across a short range using infrared and are designed to work collaboratively [18]; nevertheless, each Kilobot can be assigned a unique ID, which makes this platform suitable for testing scenarios where individual kilobots are allocated different tasks. Some of the best examples of cooperative behaviour among animals come from the world of social insects ([19],[20]) and previous research has successfully employed kilobots to simulate collective behaviour in ants ([21]). In one of these studies, a simple algorithm was designed to encompass searching behaviour by foraging kilobots to identify suitable food sources, returning to the nest to inform other nestmates about the new food source, leading them to the new food and finally bringing the food back to the colony [22]. This study provided an excellent first validation of the possibility of effectively simulating ant foraging behaviour and recruitment of nestmates using kilobots. Other studies that were published shortly after used swarm robotics to assess ant foraging, with a particular focus on the optimal distribution for collective foraging systems ([23]) or the effect of colony size on recruitment strategy ([24]). The premise of our study, instead, was based on previous work on the adaptive value of tandem communication in ants [25].

Ants are social insects that are able to communicate about profitable food and nesting sites with nestmates ([26]). One way that ants have to recruit nestmates to a known site is tandem-pair foraging, also called tandem-pair running. This strategy involves a pair of individuals, where one leads the other in pursuit of a goal, usually food or a nest site ([27]). This strategy allows for the follower to be taught how to forage or hunt [28]. Co-operation between the two members of the pair allows them to survive against other threats or to hunt more successfully. The behaviour has been seen in insect colonies and in some birds [29]. Despite its benefits, tandem running in ants is not as widespread as one would think, being mainly restricted to smaller colonies [30]. In a large colony, in fact, broadcasting information to a large number of ants at once via chemical communication (pheromones) is more efficient [31]. In addition, a colony may not use tandem-pair foraging for other reasons. It requires a form of teaching from leader to follower [32],[33] and this translates into the time that it takes for recruitment within the nest, and the lower rate of travel that tandem pairs exhibit. Furthermore, tandem running is more complex and slower than solo scouting, the more typical foraging strategy where a single and naive ant explores the surroundings for food [28], and failure results in more time lost that could be spent on foraging. In an observation of tandem-pair running [30] it was found that of 138 tandem runs 106 (76.8%) were successful with the unsuccessful runs caused by interference from vegetation, other ants, or the leader leaving the follower behind. From an engineering point of view, these two modalities of foraging in ants, each with its pro and cons, represent a powerful benchmark to test what factors influence strategy efficiency because (1) proof-of-concept is already provided by the biological system, (2) coordination of several physical tasks is needed for success, and (3) optimal behaviour requires cooperation among robots [34].

The question as to where the use of tandem running in ants is worth the extra resources that are required for its implementation was first explored by Goy et al. [25] using an agent-based simulation approach. In that study, the proposed model predicted that the spatial-temporal distribution of resources, colony size and the ratio of scouts and recruits all have a substantial impact on the success of resource procurement. The results suggested that in low resource environments, with large margins in quality and resource life-time, tandem running is favoured. In our study, we moved one step further and used these fascinating behavioural strategies displayed by ant colonies to implement a series of experiments with physical swarm robots, that were then compared with computer simulations. Past studies, in fact, have revealed how the direct comparison between real life experiments with kilobots and simulation present numerous issues, contributing to the reality gap challenge [15] [14]. These studies combined the use of real robots and simulation to reproduce both stochastic and deterministic foraging behaviours [35]. Our work is in line with this field of research, in that we used ant foraging behaviour to compare the basis for a stochastic (lone scout) vs. a deterministic (tandem running) behavioural process [25].

In our experimental design we also included the use of a pheromone trail during tandem running trails, that in nature guides the ants to return to a previously discovered food source. In swarm robotics pheromone trails have mostly been imitated by static beacon robots ([36]; [37]; [38]) that can communicate with other robots in their neighbourhood. The main advantage of this approach is that the system can be implemented with simple robots in unstructured environments; the main disadvantage is that the beacon robots are not contributing to foraging [23]. Furthermore, in our trials, we tested the effect of experimental arenas of different sizes (to reflect the variability of foraging range in the natural environment) on the success of the different foraging strategies, both with real kilobots and in simulation. Experiments were conducted in the robotics laboratory at the University of Aberdeen (see Fig.5) while the simulation were performed using the software CoppeliaSim.

## METHODS

### Laboratory Experiments with real Kilobots

An arena was used to perform experiments with real kilobots in the lab. The arena was built on a tabletop area bounded by planks of PVC, to adjust the area size as needed. To avoid the planks being nudged by the kilobots they were held in place by adhesive putty. The experiments were recorded with a Canon EOS M50 camera placed above the experimental area fig:arena. The videos were stored and the time stamps of the videos were used to determine the foraging duration. This was measured from the exact time when the kilobots started moving to the moment when the follower (in the case of a tandem pair) or the lone scout found the food. The kilobots LEDs were used to easily discern the kilobots state for accurate time keeping. The LED flashed purple before a test had commenced, displayed a solid colour during the tests, depending on the kilobots direction of movement, and turned white when the test was complete. Experiments began with a set up of kilobots reproducing the behaviour of a tandem pair (leader and follower, see Fig.3). Stationary kilobots known as beacons were positioned in a line in order to simulate a pheromone trail. Each beacon transmitted a unique ID and the tandem pair was constantly searching for a lower ID than the one being received - with food source having the lowest ID (ID 0) while every beacon in the trail had an ID that was one measure higher than the previous. Therefore, for a set-up with two beacons and a food source, the first beacon in the trail would have ID 2, while the beacon next to the food source would then have ID 1. This resulted in the tandem pair orbiting around each beacon in turn until the next beacon was detected - at which point the pair began orbiting around it. This design allowed for more beacons to be added for further experiments. The leader, follower, beacon and food kilobots all communicated with one another. All these kilobots were positioned at 8*cm* apart from each other, measured from the centre of each kilobots, with the exception of the follower that was 6*cm* from the leader. Marks on the whiteboard surface of the area were used to keep track of the initial positions and orientations of the kilobots. Once the ID for the food source was found (ID 0) the tandem leader continued its orbit until the follower was within a set distance (4.5*cm*) of the food source. At which point it was considered that the follower had found the food source and the time was recorded.

During this process, the two members of the tandem pair aimed to maintain a close distance between each other. The follower started orbiting around the beacons when it detected a distance greater than 5*cm* between itself and its leader. The leader moved when it detected a distance lower than the same threshold between itself and the follower. We tested three different scenarios, each one with a different number of beacons, corresponding to different lengths of the pheromone trail: one, two and three beacons (Fig. 2). A width of 19*cm* was chosen for the arena to ensure that the tandem pair had sufficient area to move around the beacons. The arena length instead changed according to the number of beacons used, as it had to accommodate not only the beacons themselves but also the space in between them. The smallest arena length was 25.5*cm* (one beacon), after which the arena lengths were increased by 5*cm* each time.

**Fig. 1.**
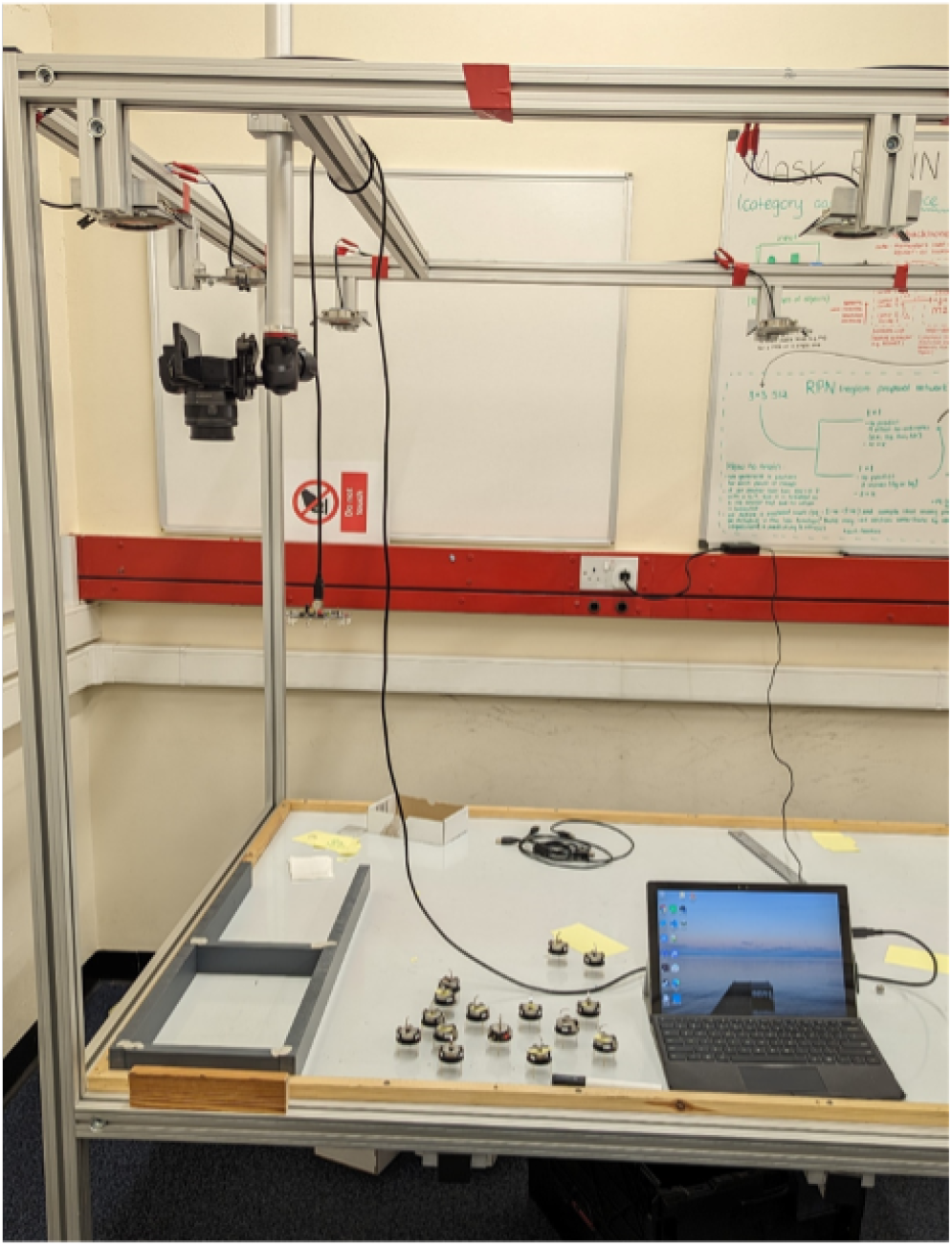
Experimental arena for Kilobots with overhead camera, kilobots, controller and laptop

**Fig. 2.**
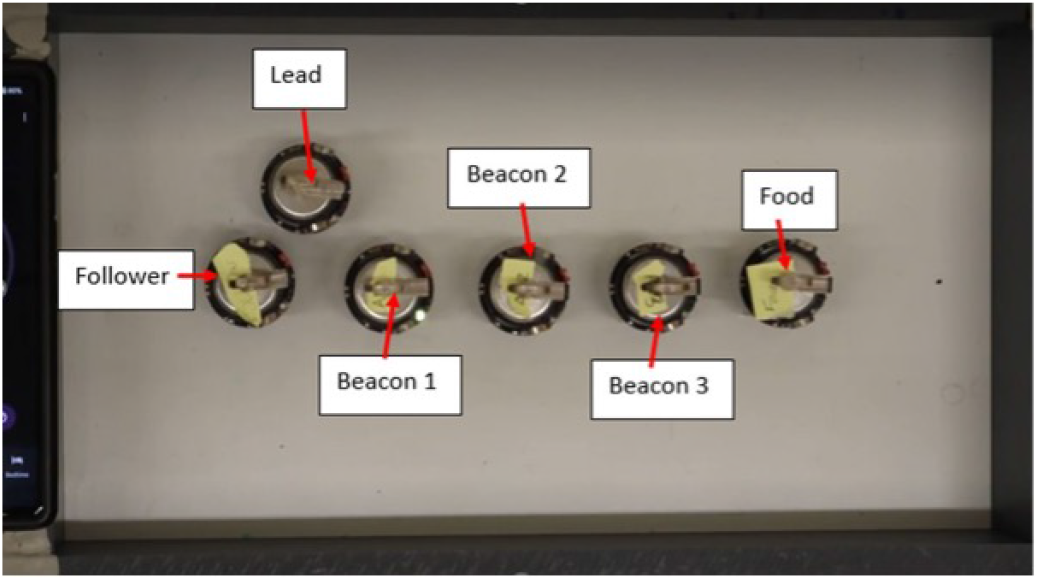
Kilobots in tandem pair configuration, the “lead” kilobot guides the “follower” using the stationary “beacons” as a guide to the “food”.

**Fig. 3.**
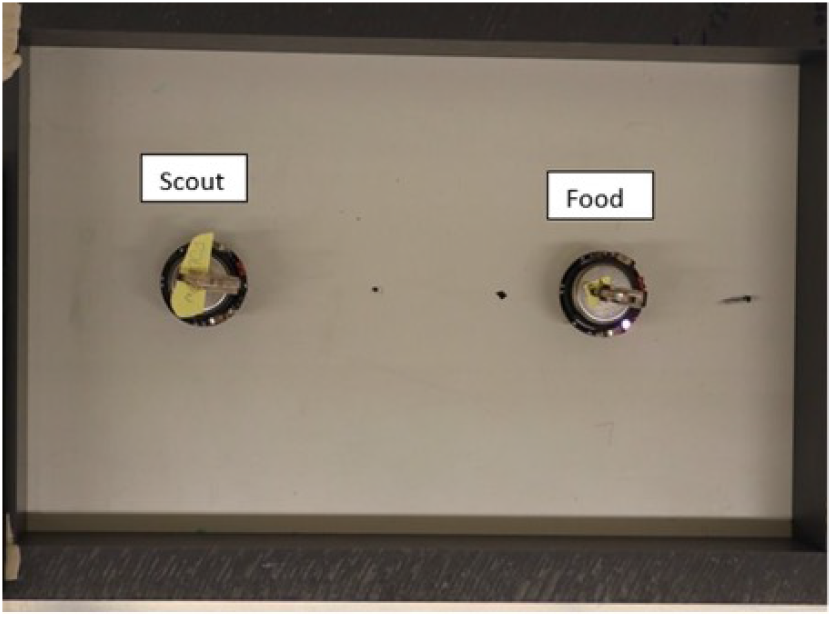
Kilobots in lone scout configuration, with lone scout (left) and food source (right).

**Fig. 4.**
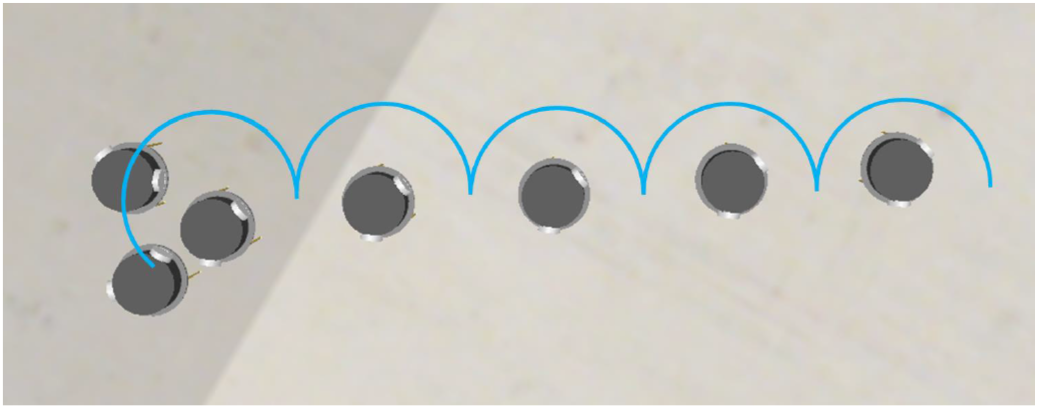
The path taken by the tandem pair kilobots following three “beacon” kilobots to the objective “food” kilobot.

**Fig. 5.**
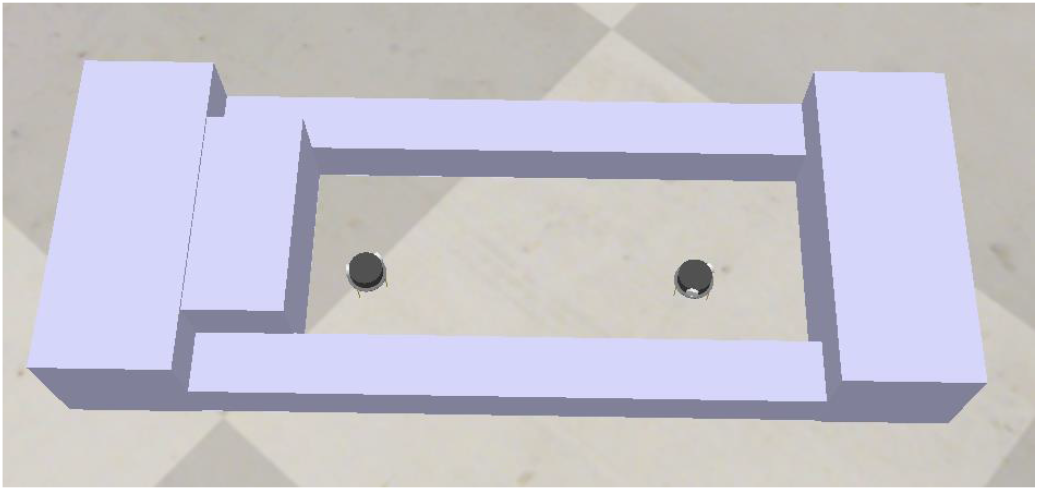
Arena for Kilobots in simulation, with lone scout (left) and food source (right).

Table 1 shows the arena size for each of the tests and the number of beacons in each arena size.

**Table 1.** Arena sizes and number of beacon for physical Kilobot experiments

| Arena Length (cm) | Arena Size (cm<sup>2</sup>) | Number of Beacons |
| --- | --- | --- |
| 25.5 | 484.5 | 1 |
| 30.5 | 579.5 | 2 |
| 35.5 | 674.5 | 3 |

The second part of the experiment involved using individual kilobots that behaved like scouts foraging for food alone. The lone scout kilobot simulated an ant scout searching for a source of food without prior knowledge of its location in the area around the nest, hence without relying on a pheromone trail. The lone scout (see Fig.3) used therefore a random search to locate profitable food sites. This was implemented by telling the lone scout kilobot to turn towards a random direction (either clockwise or anticlockwise) for 2.5 sec and then move straight for 5 sec before turning in a random direction again. To prevent a full rotation (360 degrees) at each turn, the kilobots were programmed to turn for half as much time as they moved straight. This behaviour continued until the scout came within the same set distance of the food source as described for the tandem pair. At this point the time was recorded and the experiment concluded. At the beginning of the trial, the lone scout was placed in the same position as the follower kilobot in the tandem pair as described above and the food source maintained the same position. This was done to keep the foraging distances the same for both experiments.

All kilobots were controlled and calibrated during the experiment using kilogui [39], a software that uploads programs to the kilobots *via* the overhead controller. Once the custom hex code has been uploaded, commands can be sent to the kilobots to run, stop, or reset the currently uploaded program. Calibration was done on each moving kilobot: this involved testing and adjusting values for each motor to verify that they could turn in both directions and move straight without any issues. The total number of tests completed was eight tandem-pair experiments and eight lone scout experiments.

### Simulations with CoppeliaSim

A simulation of the aforementioned robot foraging task was created with the purpose to check how transferable the observations with real kilobots would be in a virtual environment, more suitable for testing additional experimental conditions like, for example, a wider range of arena sizes.

All the behaviours designed for experiments with real Kilobots were reproduced as closely as possible in the simulation software CoppeliaSim[40] so that the outputs of the two approaches could be directly compared. For the assays with tandem pairs we utilized the same distance metrics: kilobots designated as beacons or food source were stationary and set with a unique ID corresponding to their position in the trail. The two members of the tandem pair were also set up as in the experiments with real Kilobots, with each member orbiting around the beacons on the way to the food source.

The functional CoppeliaSim kilobot model developed by K-Team [39] was utilised in the simulations with Engine Bullet 2.78: this model can simulate a multi-agent kilobot system, with capabilities that can simulate if not mimic directly the communication and locomotion of ants. While the code for the tandem pairs was an exact match to the one run on real Kilobots, the code for the lone scouts was slightly modified. As stated previously, when using the real kilobots the lone scout turned for 2.5 sec and moved straight for 5 sec. The simulated kilobots on the other hand turned and moved straight for the same period of time (5 sec). In fact, during the initial testing, it was determined that this caused the runs to fail less frequently. Simulations were run for the same arena sizes as for experiments with real kilobots, as shown in table 1.

### Combinations

In a tandem pair of foraging ants, the leader already possesses the information on where the food source is located, and acquiring that information takes time. In a set of additional tests, we decided therefore to incorporate the time required by a leader to find a new food source and added it to the time used to recruit a follower to the site. To do so, we used data obtained from the assays described above to combine a lone scout run with a tandem pair run, producing all possible combinations: this was done for both experiments with real kilobots and simulations, for all arena size. Analogously, we produced all possible combinations of two runs performed by lone scouts so that the two foraging techniques could be directly compared in a more realistic setup. This approach had two additional benefits: it allowed performing a more robust analysis on a larger set of replicates and it also mitigated against the extremely high variance that was observed in the duration of individual runs performed by lone scouts, both in the assays with real kilobots and in the simulations.

### Statistical Analyses

We used SPSS Statistics software to analyze data from experiments with real kilobots, simulations with CoppeliaSim and combinations of tandem pair and random search runs performed by lone scouts. Independent-samples T tests were performed to determine whether the duration of runs for tandem pairs was significantly different from random searches of lone scouts for each arena size - or to test whether the duration of a tandem pair + a random search was significantly different from two random searches in the combination assays.

## RESULTS

Experiments with real kilobots revealed that the duration of the search to find the ”food” beacon between tandem pairs and lone scouts was very similar in the arena with the smallest size (484.5*cm*^2^, Table 2): The difference between tandem pair (M = 42.280 sec, SD = 8.629) and lone scouts (M = 52.687 sec, SD = 46.297) was not significant (t(18) = -0.466, p = 0.324). Starting from the arena of intermediate size (579.5*cm*^2^) a trend was observed for tandem pair strategy (M= 86.660 sec, SD = 23.363) to become faster than the lone scout strategy (M = 112.793 sec, SD = 88.149), though the difference was not significant (t(18) = -0.645, p = 0.264). The trials in the largest arena (674.5*cm*^2^) revealed the most evident difference in duration, with tandem pairs (M = 129.440 sec, SD = 22.868) finding the food site significantly quicker than lone scouts (M = 180.007 sec, SD = 104.229; t(17) = -1.756, p = 0.48).

**Table 2.** The mean time (seconds) and standard deviation for each category of kilobot experiment.

|  |  | Arena Size |  |  |
| --- | --- | --- | --- | --- |
|  |  | 484.5 cm<sup>2</sup> | 579.5 cm<sup>2</sup> | 674.5 cm<sup>2</sup> |
| Experimental | Tandem Pair | 42.820±8.629 | 86.660±23.363 | 129.440±22.868 |
|  | Lone Scout | 52.687±46.297 | 112.793±88.149 | 180.007±104.229 |
| Simulation | Tandem Pair | 155.595±9.665 | 265.187±9.038 | 329.072±21.881 |
|  | Lone Scout | 153.375±145.852 | 220.449±162.970 | 256.412±113.550 |
| Experimental Combination | Lone Scout + Tandem Pair | 95.513±16.916 | 199.453±32.193 | 309.447±38.065 |
|  | Lone Scout + Lone Scout | 105.387±16.916 | 225.587±32.193 | 360.013±38.065 |
| Simulation Combination | Lone Scout + Tandem Pair | 308.970±53.266 | 485.637±59.518 | 585.484±41.470 |
|  | Lone Scout + Lone Scout | 306.750±53.266 | 440.899±59.518 | 512.824±41.470 |

Simulations with CoppeliaSim revealed that, similarly to real kilobots, there was no difference in the average amount of time spent by tandem pairs and lone scouts to locate the food in the smallest arena size: for tandem pairs (M = 155.595 sec, SD = 9.665) and for lone scouts (M = 153.375 sec, SD = 145.851) the difference was not statistically significant (t(14.363) = 0.059, p = 0.477). It is interesting to note that the average time to find the food in the smallest arena was three times higher in the simulations that in the experiments with real kilobots. Starting from the arena of intermediate size it was possible to see a difference in the time needed to find the food within the two foraging strategies: the trend, however, was opposite to what reported for experiments with real kilobots, with tandem pairs (M = 265.187 sec, SD = 9.038) taking longer time than lone scouts (M = 220.499 sec, SD = 162.971, t(14.255) = 1.058, p = 0.154). This trend was even more visible in the arena of largest size, where the difference was significant between tandem pairs (M= 329.072 sec, SD = 21.881) and lone scouts (M = 256.412 sec, SD = 113.550, t(16.573) = 2.351, p = 0.16).

Combination tests produced very similar outputs to what observed in data from the initial experiments. The difference in searching times for a tandem pair + a random search by a lone scout *vs*. two random searches by lone scouts was minimal for the smallest arenas, while it became progressively larger for arenas of intermediate and largest size (Fig. 6, Fig.7 and Table 2). As observed for the initial experiments, tandem pair strategy became progressively faster than lone scout search for real kilobots as arena size increased, while in simulations lone scout strategy became faster. Differently from the output of the initial experiments, all comparisons for real kilobot combinations were statistically different (*p <* 0.001) while in the case of simulations, only arenas of intermediate and largest size showed significantly different search times (*p <* 0.001) while this was not the case for the arena of smallest size (p = 0.53).

**Fig. 6.**
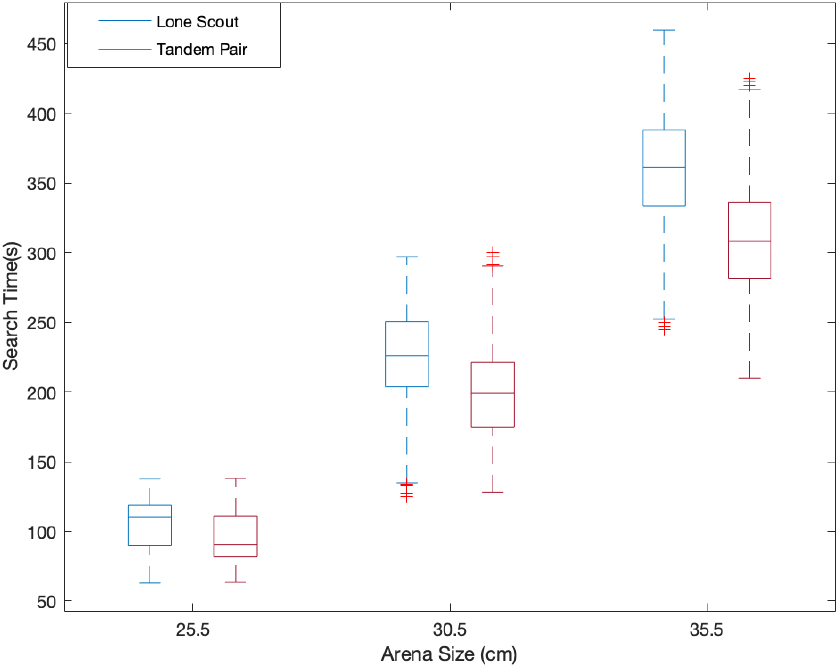
The time elapsed for the combinations of the real-life experiments, for each arena size

**Fig. 7.**
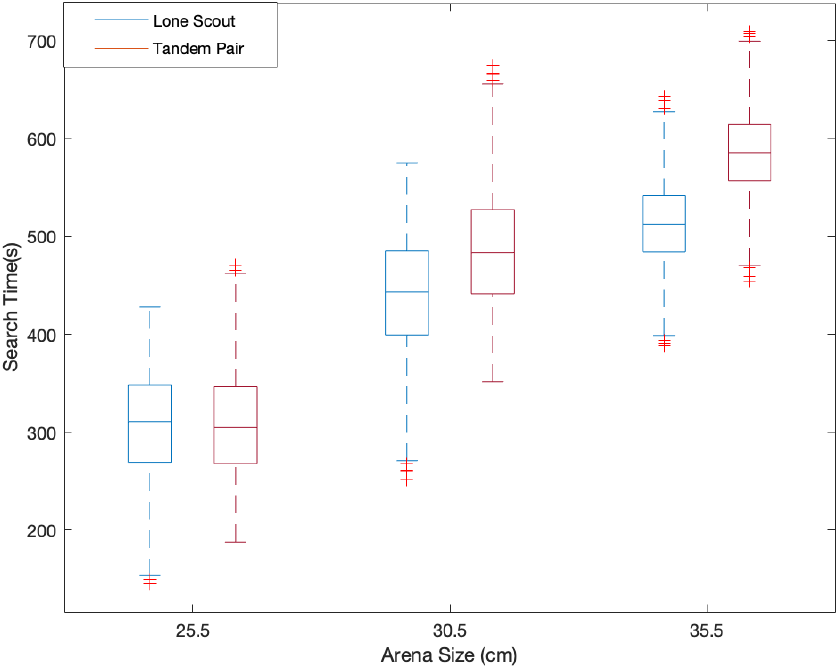
The time elapsed for the combinations of the simulated experiments in CoppeliaSim, for each arena size

Table 2 shows the results obtained for the experiments performed with the real and simulated Kilobots.

## DISCUSSION

In this study we aimed to use kilobots to better characterize two foraging strategies that are typical in ant colonies: tandem pair recruitment and lone scout foraging. We compared the amount of time that was necessary to find a food site for kilobots adopting one of the two strategies in experimental arenas of increasing size - to simulate increasing distance of the food source from the colony - and we compared these scenarios using real kilobots in the lab *vs*. using simulations with virtual kilobots using CoppeliaSim. Similarly, for the two approaches, there was no difference between tandem pairs and lone scouts for the smallest arenas (though time spent in simulations was visibly higher than with real kilobots) while differences between the two strategies started to emerge from intermediate arenas and became more prominent for the arenas of largest sizes - i.e., when the food source was the farthest from the foraging ants. These differences, however, followed opposite trends and these observations were confirmed when performing combination tests to control for initial search time of leaders in tandem pairs and for the high variation observed across individual random searches. Our results highlight how difficult it is to use simulations to model the behaviour of kilobots in real life - something that has been reported before in the literature [15], [16]. We discuss these results in the context of ant foraging behaviour as well as in relation to the reality gap in robotics and other applications of kilobots and robotics to biological scenarios.

In Goy et al. the authors found that the time needed for recruits and scouts (equivalent to our tandem pairs and lone scouts, respectively) to find the food was highly affected by the combination of conditions in the experimental environment, namely colony size, quantity and quality of the food and distance from the nest. In our study, the number of variables assessed was limited, as we wanted to focus on the comparison between experiments with real kilobots *vs*. simulations: we tested increasing arena sizes (hence distance of the food from the colony), while we had only one food item of high quality. Therefore, a direct comparison with Goy et al. is not straightforward but it is still noteworthy that our experiment with real kilobots aligned well with what reported by the authors for limited numbers of high quality food sources: in this scenario, tandem running is faster than the random search of a lone scout. It will be interesting to assess in the future the output of experiments with real kilobots when adding more variable like, for example, more food items of different value. As for the results of our simulations, the patterns observed going in opposite direction compared to real kilobots and simulations performed in Goy et al., are difficult to explain. One obvious reason could be that lone scouts in simulated environment are extremely fast - due to the lack of physical constraints in a virtual experimental arena - and therefore they easily out-compete tandem pairs in all scenarios. This explanation, however, is defeated by the fact that in all simulations kilobots took visibly longer time to complete the search compared to real kilobots (up to three times longer for the smallest arenas) no matter what the strategy adopted was. The source of this inconsistency, therefore, remains unsolved and invites for more assays to compare simulations with real experiments and fully address the long-standing reality gap, perhaps increasing the arena size even further to test whether the same trends are confirmed.

When it comes to assessing our results in the context of foraging behaviour in ant colonies similar considerations as above still hold: in fact, previous studies reported that tandem followers are faster than lone scouts to find a single food source [28]. This is corroborated by swarm robotics research showing that recruitment significantly increased foraging efficiency compared with experiments where robots could not communicate [41]. It was also found that ants were more likely to perform tandem runs and they travelled faster when the food source was more distant [30] - similar to our arenas of increasing size. In this last study, however, the authors also report that tandem running has costs, mainly associated with the logistic challenges of two ants moving together in a complex environment, characterized by a wide range of obstacles such as impervious substrates and predators (just to mention a few), and with the possibility of breakouts in the tandem pair. For example, the previous literature reported that in different species of ants, individual foragers can be 1.5 to 4 times faster foraging than tandem pairs [42], [28]; furthermore, a breakup of the tandem-pair was reported 32 times out of 138 observed runs (or 23.2%) in [30]. In a separate set of experiments with real kilobots, we observed that three out of the eight tests with tandem pairs failed, which means a failure rate of 37.5% (*unpublished data*). This might have been caused by localisation errors, for example their infrared proximity sensors malfunctioning. This seems to be a much higher rate than what observed in nature, but we must take into account that the sample size in our experiments was limited and this might have influenced the outcome. Using augmented reality for Kilobots (ARK) [43], [23], which provides Kilobots with extra information based on their location and state, could reduce the errors that occurred during our experiments and likely caused the occurrence of the reported breakouts.

CoppeliaSim was chosen as the simulation environment due to its various physics engines and more efficient CPU use, compared to other simulators [44]. This study clearly show that the CoppeliaSim platform is not able to accurately simulate the complexity of real-life experiments with kilobots. One of the most striking examples of this discrepancy was observed for the orbiting functionality, which was not performed by the kilobots in the real experiment as it was in the simulation [16]. The reality-gap is a recognised and limiting phenomenon that has significant implications in the field of bio-inspired robotics. Simulations are fundamental to inform planning experiments with real kilobots but, at the same time, cannot fully capture the complexity of the environment a robot is acting in - which is nevertheless significantly simpler than the real world where organisms live and behave. Even in the super-simplified foraging-arena of a robotic lab there are many factors such as individual mass or collisions and traction among robots [14] that might have an influence on the outcome of the robot-robot interactions, but are only partially considered by the simulation software [15]. Some help could come from more advanced physics engines, the programs that simulate physics phenomena or carefully selecting among the available physics engines available in a simulator. A recent study, in fact, reported that Open Dynamics Engine or ODE provides results that are the closest to the output of a real environment among all tested physics engines [15].

Our study introduces an innovative approach in the comparative assessment of complex animal behaviours using kilobots: however, it only represents a first step that paves the way to a range of possible implementations in the future, aimed not only to producing a more robust system, but also one that more closely resembles the reality of the behaviour of interest. One obvious development of this research will be to test whether the same patterns that we observed (i.e., faster foraging times for tandem-pair strategy compared to lone scouts) hold for arena of larger sizes. It would be more convenient to test this in simulations of course, as less challenging logistically; however, our findings implicate that this will not be enough and experiments with real kilobots will also be needed. Secondly, in this series of experiments, we did not exploit that full potential of the kilobots platform, designed so that a large number of agents can act in parallel as a swarm: this was motivated by the need to keep the experimental design simple, being this the first series of trials of this kind. In the future it will be interesting to exploit the ability of swarm robotics to successfully carry out a task by using a high number of agents. This will translate in multiple kilobot ants performing the food search in parallel, adopting both tandem pair and lone scout strategies, which more closely reflects the dynamics occurring within an ant colony. Finally, another aspect that could be implemented in the future is the presence of competing food qualities, something that is also recurring in real-life biological systems: ants - similarly to all other foraging organisms - continuously investigate the environment to detect the most rewarding food sources and update their feeding strategy accordingly.

### Conclusions

The main goal of this project was to establish a framework for developing biologically inspired models using Kilobots. We were able to successfully simulate ant foraging behaviour in a controlled environment, establishing a platform that can be used in the future to study and simulate other complex animal behaviours. Our findings highlight the need to fill the reality gap, so that simulation can be more reliable in informing real-life experiments, and also the power of Kilobots to provide an experimental platform where it is possible to compare different biological phenomena where multiple agents work in parallel. Immediate application of our findings could be, for example, in the fields of swarm intelligence, collective decision making and navigation techniques and strategies. The potential application of these findings to more distant fields such as logistics, agriculture, and search and rescue highlights the importance of this research and its findings.

## Acknowledgments

We thank the Schools of Biological Sciences and Engineering for their support. We thank Bruce McKay for his contribution to the preliminary simulation work.

## Author contributions statement

F.M, M.E.G. and N.E. conceived the experiments; N.E. and I.Y. wrote the code for the real-life and simulation experiments; A.C. conducted the real-life experiments while N.E. performed the simulation trials; J.K. analysed the results. All authors contributed to the preparation of the first draft and reviewed the manuscript.

## Previous presentation

These results were not previously presented at any conference.

## Preprints

A preprint of this article is published at [DOI].

## Data availability

The code and videos underlying this article are available in ”Testing the reality gap with kilobots performing two ant-inspired foraging behaviours” - code and videos at https://github.com/Niamh-Ellis/Kilobot-Ant-Code/tree/main - and can be accessed with https://doi.org/10.5281/zenodo.11410123.

## References

1. Rory Cellan-Jones. Robots ‘to replace up to 20 million factory jobs’ by 2030. BBC News.

2. Chuck Thorpe and Hugh Durrant-Whyte. Field robots. In Robotics Research: The Tenth International Symposium, pages 329–340. Springer, 2003.

3. Barbara Webb. What does robotics offer animal behaviour? Animal behaviour, 60(5):545–558, 2000.

4. Auke J Ijspeert. Biorobotics: Using robots to emulate and investigate agile locomotion. science, 346(6206):196–203, 2014.

5. Manuele Brambilla, Eliseo Ferrante, Mauro Birattari, and Marco Dorigo. Swarm robotics: a review from the swarm engineering perspective. Swarm Intelligence, 7:1–41, 2013.

6. Guang-Zhong Yang, Jim Bellingham, Pierre E Dupont, Peer Fischer, Luciano Floridi, Robert Full, Neil Jacobstein, Vijay Kumar, Marcia McNutt, Robert Merrifield, et al. The grand challenges of science robotics. Science robotics, 3(14):eaar7650, 2018.

7. Christopher M Cianci, Xavier Raemy, Jim Pugh, and Alcherio Martinoli. Communication in a swarm of miniature robots: The e-puck as an educational tool for swarm robotics. In International Workshop on Swarm Robotics, pages 103–115. Springer, 2006.

8. Jiangfan Yu, Lidong Yang, and Li Zhang. Pattern generation and motion control of a vortex-like paramagnetic nanoparticle swarm. The International Journal of Robotics Research, 37(8):912–930, 2018.

9. Merihan Alhafnawi, Edmund R Hunt, Severin Lemaignan, Paul O’Dowd, and Sabine Hauert. Mosaix: a swarm of robot tiles for social human-swarm interaction. In 2022 International Conference on Robotics and Automation (ICRA), pages 6882–6888. IEEE, 2022.

10. Florian Berlinger, Melvin Gauci, and Radhika Nagpal. Implicit coordination for 3d underwater collective behaviors in a fish-inspired robot swarm. Science Robotics, 6(50), 2021.

11. Nick Jakobi, Phil Husbands, and Inman Harvey. Noise and the reality gap: The use of simulation in evolutionary robotics. In Advances in Artificial Life: Third European Conference on Artificial Life Granada, Spain, June 4–6, 1995 Proceedings 3, pages 704–720. Springer, 1995.

12. Antoine Ligot and Mauro Birattari. On mimicking the effects of the reality gap with simulation-only experiments. In Swarm Intelligence: 11th International Conference, ANTS 2018, Rome, Italy, October 29–31, 2018, Proceedings 11, pages 109–122. Springer, 2018.

13. Antoine Ligot and Mauro Birattari. Simulation-only experiments to mimic the effects of the reality gap in the automatic design of robot swarms. Swarm Intelligence, 14(1):1–24, 2020.

14. Paul Christiano, Zain Shah, Igor Mordatch, Jonas Schneider, Trevor Blackwell, Joshua Tobin, Pieter Abbeel, and Wojciech Zaremba. Transfer from simulation to real world through learning deep inverse dynamics model. arXiv preprint 1610.03518, 2016.

15. Andreas Meier, Sascha Carroccio, Rolf Dornberger, and Thomas Hanne. Discussing the reality gap by comparing physics engines in kilobot simulations. Journal of Robotics and Control (JRC), 2(5):441–447, 2021.

16. Vivienne Jia Zhong, Rolf Dornberger, and Thomas Hanne. Comparison of the behavior of swarm robots with their computer simulations applying target-searching algorithms. International Journal of Mechanical Engineering and Robotics Research, 7(5), 2018.

17. Michael Rubenstein, Christian Ahler, Nick Hoff, Adrian Cabrera, and Radhika Nagpal. Kilobot: A low cost robot with scalable operations designed for collective behaviors. Robotics and Autonomous Systems, 62(7):966–975, 2014. Reconfigurable Modular Robotics.

18. S. Prasath Kumar, A. Ravindiran, S. Meganathan, N. Oral Roberts, and N. Anbarasi. Swarm robot materials handling paradigm for solar energy conservation. Materials Today: Proceedings, 46:3924–3928, 2021. International Conference on Materials, Manufacturing and Mechanical Engineering for Sustainable Developments-2020 (ICMSD 2020).

19. Elva J. H. Robinson and Jessica L. Barker. Inter-group cooperation in humans and other animals. Biology Letters, 13(3):20160793, 2017.

20. António M. M. Rodrigues, Jessica L. Barker, and Elva J. H. Robinson. From inter-group conflict to inter-group cooperation: insights from social insects. Philosophical Transactions of the Royal Society B: Biological Sciences, 377(1851):20210466, 2022.

21. Michael Rubenstein, Alejandro Cornejo, and Radhika Nagpal. Programmable self-assembly in a thousand-robot swarm. Science, 345(6198):795–799, 2014.

22. Akimasa Otsuka, Ryota Ueda, and Fusaomi Nagata. Experiment of imitating ant feeding behavior using kilobot. In 2017 IEEE International Conference on Mechatronics and Automation (ICMA), pages 531–535, 2017.

23. Mohamed S. Talamali, Thomas Bose, Matthew Haire, Xu Xu, James A. R. Marshall, and Andreagiovanni Reina. Sophisticated collective foraging with minimalist agents: a swarm robotics test. Swarm Intelligence, 14(1):25–56, Mar 2020.

24. Siddharth Mayya, Pietro Pierpaoli, and Magnus Egerstedt. Voluntary retreat for decentralized interference reduction in robot swarms. In 2019 international conference on robotics and automation (ICRA), pages 9667–9673. IEEE, 2019.

25. Natascha Goy, Simone M. Glaser, and Christoph Grüter. The adaptive value of tandem communication in ants: Insights from an agent-based model. Journal of Theoretical Biology, 526:110762, 2021.

26. Elva J.H. Robinson, Francis L.W. Ratnieks, and M. Holcombe. An agent-based model to investigate the roles of attractive and repellent pheromones in ant decision making during foraging. Journal of Theoretical Biology, 255(2):250–258, 2008.

27. E.L. Franklin. The journey of tandem running: the twists, turns and what we have learned. Insectes Sociaux, 61:1–8, 2014.

28. Nigel Franks and Thomas Richardson. Teaching in tandem-running ants. Nature, 439:153, 02 2006.

29. Martin J Folk. Cooperative hunting of avian prey by a pair of Bald Eagles. Florida field naturalist, 1992.

30. C. Grüter, M. Wüst, A. P. Cipriano, and F. S. Nascimento. Tandem recruitment and foraging in the ponerine ant pachycondyla harpax (fabricius). Neotropical Entomology, 47(6):742–749, Dec 2018.

31. Sophie EF Evison, Owen L Petchey, Andrew P Beckerman, and Francis LW Ratnieks. Combined use of pheromone trails and visual landmarks by the common garden ant lasius niger. Behavioral Ecology and Sociobiology, 63:261–267, 2008.

32. William J.E. Hoppitt, Gillian R. Brown, Rachel Kendal, Luke Rendell, Alex Thorton, Mike M. Webster, and Kevin N. Laland. Lessons from animal teaching. Trends in ecology & evolution, 23(9):486–493, 2008.

33. Takao Sasaki, Leo Danczak, Beth Thompson, Trisha Morshed, and Stephen C Pratt. Route learning during tandem running in the rock ant temnothorax albipennis. Journal of Experimental Biology, 223(9):jeb221408, 2020.

34. Alan FT Winfield. Foraging robots. Encyclopedia of complexity and systems science, 6:3682–3700, 2009.

35. Qi Lu, Antonio D Griego, G Matthew Fricke, and Melanie E Moses. Comparing physical and simulated performance of a deterministic and a bio-inspired stochastic foraging strategy for robot swarms. In 2019 International Conference on Robotics and Automation (ICRA), pages 9285–9291. IEEE, 2019.

36. Barry Brian Werger and Maja J Mataric. Robotic food chains: Externalization of state and program for minimal-agent foraging. 1996.

37. David Payton, Mike Daily, Regina Estowski, Mike Howard, and Craig Lee. Pheromone robotics. Autonomous Robots, 11:319–324, 2001.

38. Shervin Nouyan, Roderich Groß, Michael Bonani, Francesco Mondada, and Marco Dorigo. Teamwork in self-organized robot colonies. IEEE Transactions on Evolutionary Computation, 13(4):695–711, 2009.

39. Julien Tharin. Kilobot user manual, Jan 2016.

40. Coppeliasim robotics. https://www.coppeliarobotics.com/. Accessed: 21st October 2023.

41. Michael JB Krieger, Jean-Bernard Billeter, and Laurent Keller. Ant-like task allocation and recruitment in cooperative robots. Nature, 406(6799):992–995, 2000.

42. Patrick Schultheiss, Chloé A Raderschall, and Ajay Narendra. Follower ants in a tandem pair are not always naïve. Scientific reports, 5(1):10747, 2015.

43. Andreagiovanni Reina, Alex J Cope, Eleftherios Nikolaidis, James AR Marshall, and Chelsea Sabo. Ark: Augmented reality for kilobots. IEEE Robotics and Automation letters, 2(3):1755–1761, 2017.

44. Lenka Pitonakova, Manuel Giuliani, Anthony Pipe, and Alan Winfield. Feature and performance comparison of the v-rep, gazebo and argos robot simulators. In Towards Autonomous Robotic Systems: 19th Annual Conference, TAROS 2018, Bristol, UK July 25-27, 2018, Proceedings 19, pages 357–368. Springer, 2018.

